# abmR: an R package for agent-based model analysis of large-scale movements across taxa

**DOI:** 10.1101/2021.09.15.460374

**Authors:** Benjamin Gochanour, Javier Fernández-López, Andrea Contina

## Abstract

1. Agent-based modeling (ABM) shows promise for animal movement studies. However, a robust, open-source, and spatially explicit ABM coding platform is currently lacking.
2. We present abmR, an R package for conducting continental-scale ABM simulations across animal taxa. The package features two movement functions, each of which relies on the Ornstein-Uhlenbeck (OU) process.
3. The theoretical background for abmR is discussed and the main functionalities are illustrated using example populations.
4. Potential future additions to this open-source package may include the ability to specify multiple environmental variables or to model interactions between agents. Additionally, updates may offer opportunities for disease ecology and integration with other R movement modeling packages.

## 1. INTRODUCTION

Animal movement is a complex behavioral trait that affects the survival of populations and species across taxa (Berg, 1983; Dingle, 2014). Long- and short-distance movements can follow predictable environmental constraints, allowing populations to take advantage of seasonal food resources (e.g., migration), or more opportunistic, such as in the case of dispersal behaviors aimed at avoiding predators or finding potential mates (Giuggioli & Bartumeus, 2010). Thus, wild animals make decisions often based on environmental cues that lead to movement patterns characteristic of different populations across the landscape (Nathan et al., 2008; Dodge et al., 2014). However, obtaining a comprehensive understanding of large-scale animal movement behavior and population occurrence under climate change scenarios or habitat loss has proven to be a challenge (Araujo & Guisan, 2006). Moreover, while the research toolbox in movement ecology studies has seen a considerable expansion over the last two decades due to technological advancements of the tracking devices and molecular markers (Cushman and Lewis, 2010; Williams et al., 2020), the limitation of scaling up individual data to population-level inferences is still a substantial obstacle (Hawkes, 2009; but see Holdo & Roach, 2013). A promising research approach that may overcome the limitations of wildlife movement studies hindered by small sample sizes is represented by computer simulations within an Agent-based Modeling (ABM) framework (Tang & Bennett, 2010; Bridge et al., 2017).

The core principle of ABM is to simulate a set of entities, called agents, which are defined by intrinsic properties as well as behavioral rules governing their interactions with the environment (Grimm & Railsback, 2013). That is, agents are described by their inherent attributes while dynamically interacting with external conditions such as the co-occurrence of other agents and/or changing features of their environmental setting. Thus, ABM is used in many areas, including biology, disease risk, social sciences, and economics (Polhill et al., 2008; Grimm & Railsback, 2013; Kilmek et al., 2015; Willem et al., 2017) with the unifying goal of investigating and predicting the dynamics of complex systems (Grimm et al., 2005). In particular, wildlife studies have adopted the ABM approach to simulate population growth, reproduction, mortality rate, energy budget, and migration ecology, just to cite a few (Brown & Robinson, 2006; Lustig et al., 2019; Aurbach et al., 2020; Goldstein et al., 2021). However, we currently lack a robust and spatially explicit ABM coding platform for the implementation of large-scale animal movement investigations (but see Thiele et al. (2012) and Chubaty and McIntire (2021)). Here we present a novel ABM framework in R programming language (R Core Team, 2020) for applications in animal behavior and movement ecology.

## 2. PACKAGE OVERVIEW

abmR allows for both computation and visualization of agent movement trajectories through a set of behavioral rules based on environmental parameters. The two movement functions, `moveSIM` and `energySIM`, provide the central functionality of the package, allowing the user to run simulations using an Ornstein-Uhlenbeck movement model (Uhlenbeck & Ornstein, 1930; hereafter OU). Additional functions provide a suite of visualization and data summarization tools intended to reduce the effort needed to go from results to presentation-ready figures and tables (Table 1). The package is currently available on the Comprehensive R Archive Network (CRAN).

**Table 1.** Functions contained in the abmR package (v. 1.0.6). For more complete function descriptions, consult the abmR manual.

| Function | Usage |
| --- | --- |
| `moveSIM` | Runs agent-based model movement simulations based on environmental data |
| `moveVIZ` | Creates a plot or table of `moveSIM()` results |
| `energySIM` | Runs agent-based model movement and energy budget simulations based on environmental data |
| `energyVIZ` | Creates a plot or table of `energySIM()` results |
| `tidy_results` | Prints results from `moveSIM()` or `energySIM()` in an easier-to-read table |
| `get_ex_data` | Downloads data that is used in examples in vignette and documentation |
| `as.species` | Creates object of class 'species' for input into `moveSIM()` or `energySIM()` |

The abmR package is built to facilitate customization of movement model parameters within the R environment (Fig. 1). Parameters affecting agent behavior can be obtained from real-world observations (e.g., GPS data) or approximated from the literature and then manually entered as arguments into abmR functions. Thus, if the user knows beforehand the behavior and the ecological constraints of the agents, such as land-cover preferences (e.g., vegetation composition and structure) or movement direction and average distance traveled per day, they may transfer this knowledge into abmR to study changes in energy consumption, mortality rate or alternative routes across different environmental conditions (e.g., raster layers). Alternatively, for simulations of ‘synthetic species’ designed to compare relative mortality rates across habitats, for example, the user may develop an analytical framework within abmR without exploring the ecology of a particular organism. In this case, the movement parameters may not be realistic, but selected with the unique goal of comparing a set of scenarios (e.g., movements in predicted future land-use changes) while studying the emerging properties of the system (Yin et al. 2022). Finally, abmR functionalities can be integrated with other approaches, such as a step-selection function (Thurfjell et al. 2014) and/or Bayesian statistics to estimate movement parameters and explore migration patterns or habitat occupancy of a particular set of agents (Joo et al. 2020; Cullen et al. 2022).

**Figure 1.**
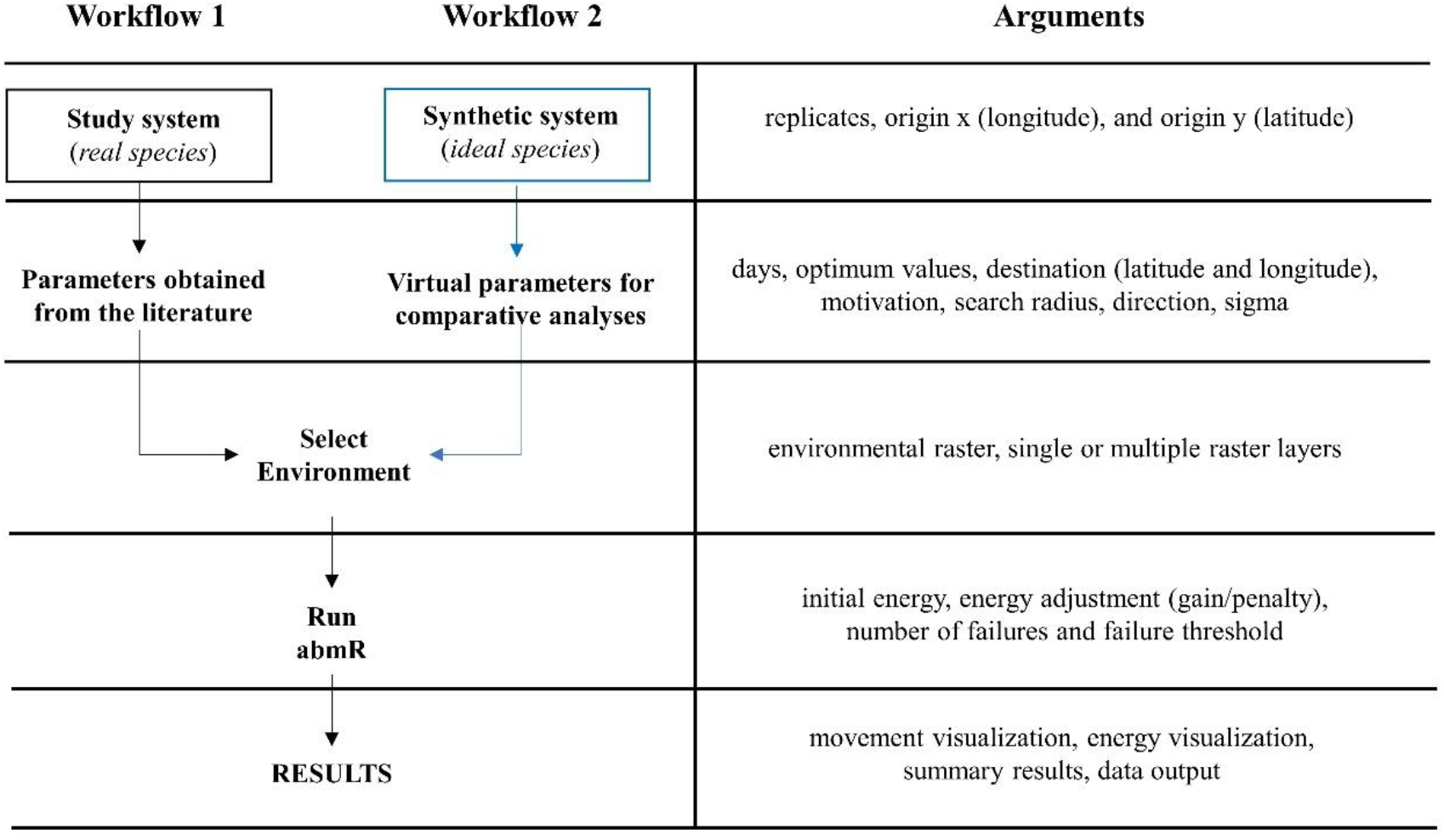
abmR workflow from data input to result exploration. The black box and black arrows (Workflow 1, left) show the sequence of the analytical steps, starting with the identification of the study system (species or population), followed by the acquisition of the movement parameters from previous observations or the literature. After selecting the environmental conditions (e.g., raster layers), users can run a suite of abmR functions to study movements, changes in energy consumption or mortality rate. The blue box and blue arrows (Workflow 2, right) show an alternative analytical workflow in which users can create synthetic species for comparative analyses across predicted ecological and environmental scenarios (e.g., loss of migratory behavior and habitat fragmentation). The panel to the right of both workflows shows a list of the relative arguments that can be entered in abmR across each step of the analysis.

Both movement functions used by abmR rely on the same OU approach. The OU process is described by the following stochastic differential equation:

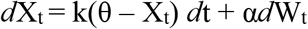

where W_t_ is a standard Brownian motion on t ∈ [0, ∞), k is a positive constant that controls the rate of reversion to the long-term mean, θ, of the OU process (see Blomberg et al 2020), and α > 0 is a constant controlling the volatility of the Brownian motion. In abmR, given the current agent location (x_t_, y_t_), agent location at the subsequent timestep (x_t+1, yt+1_) is modeled according to the following equations:

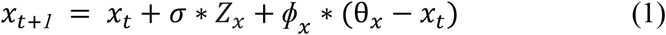

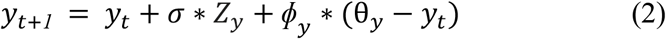

Here, *σ* is a user-specified multiplier on the random terms Z_x_ and Z_y_, two numbers drawn from the Normal (0,1) distribution. In addition, *ϕ_x_* and *ϕ_y_* are movement motivation or attraction strength for the OU process in the longitude and latitude coordinates, respectively, while θ_*x*_ and θ_*y*_ are optimal x (longitude) and y (latitude) coordinates, respectively. It is assumed that the origin point (x_1_, y_1_) is known. The OU model given in (1) and (2) performs similarly to a spring-coil (Fig. 2). Greater distance from optimal coordinates θ_*x*_ and θ_*y*_ acts like a compressed spring to propel distant agents towards θ_*x*_ and θ_*y*_. On the other hand, agents closer to θ_*x*_ and θ_*y*_ will travel a shorter distance on that timestep. However, the length of movement also depends on *ϕ_x_* and *ϕ_y_*, because these motivations serve as a multiplier on (θ_*x*_ – *x_t_*) and (θ_*y*_ – *y_t_*), respectively (Eqns. 1 and 2).

**Figure 2.**
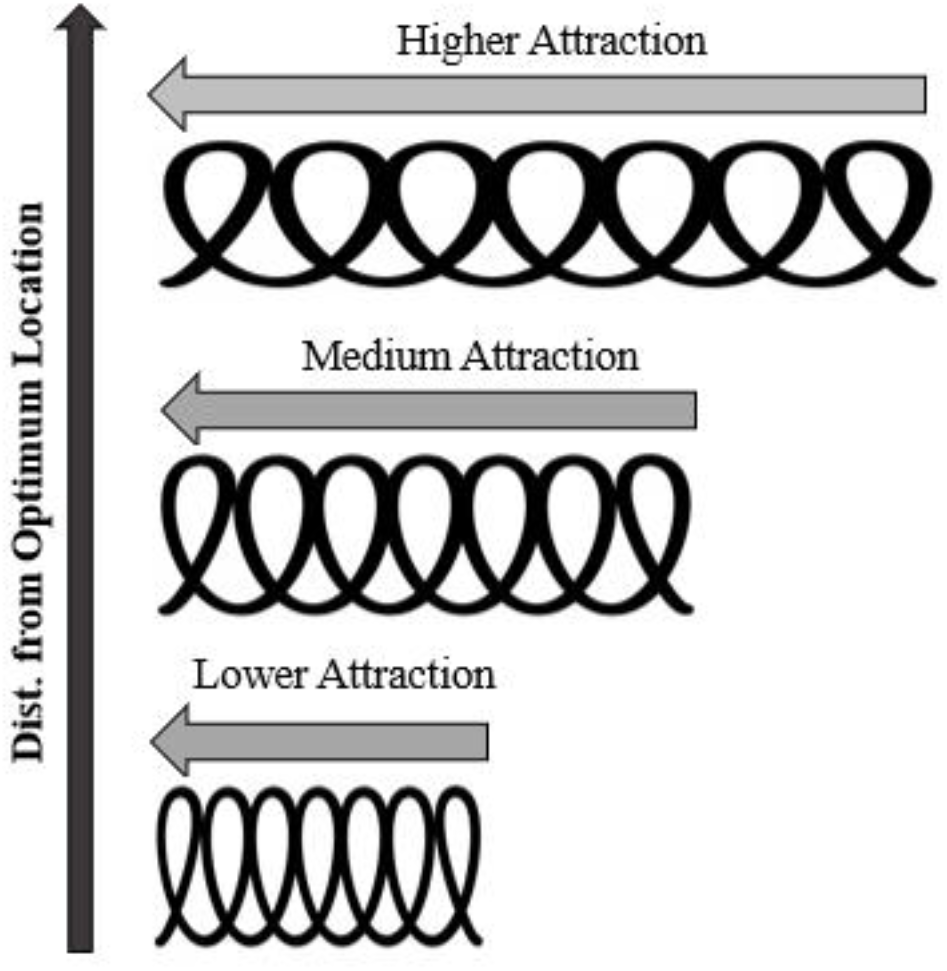
The Ornstein-Uhlenbeck (OU) model given in (1) and (2) performs like a spring-coil: agents further from their target location experience higher attraction (and travel further), while agents closer to their destination experience lesser attraction (and travel less far).

While the two movement functions are distinct (see below), each follows the same basic two steps. The first, large-scale searching, is illustrated in Fig. 3 and follows an elliptical pattern, as opposed to a circular distribution, to account for directional migratory impulses (e.g., genetically controlled migratory behaviors or seasonal Zugunruhe; Merlin & Liedvogel, 2019). This step finds the ‘optimum’ location for each agent (θ_*x*_ and θ_*y*_ coordinates from Eqns. (1) and (2)). The coordinates θ_*x*_ and θ_*y*_ are determined according to an algorithm that selects the location whose observed environmental value has the least possible difference from the user-specified optimal raster value. For `moveSIM`, this user-specified optimal raster value is supplied directly, while for `energySIM` it is the average of the lower and upper bounds of a user-specified optimum range. The optimum value or range of values specified depends on the modeling scenario and the type of environmental raster that is used (e.g., vegetation, temperature, etc.).

**Figure 3.**
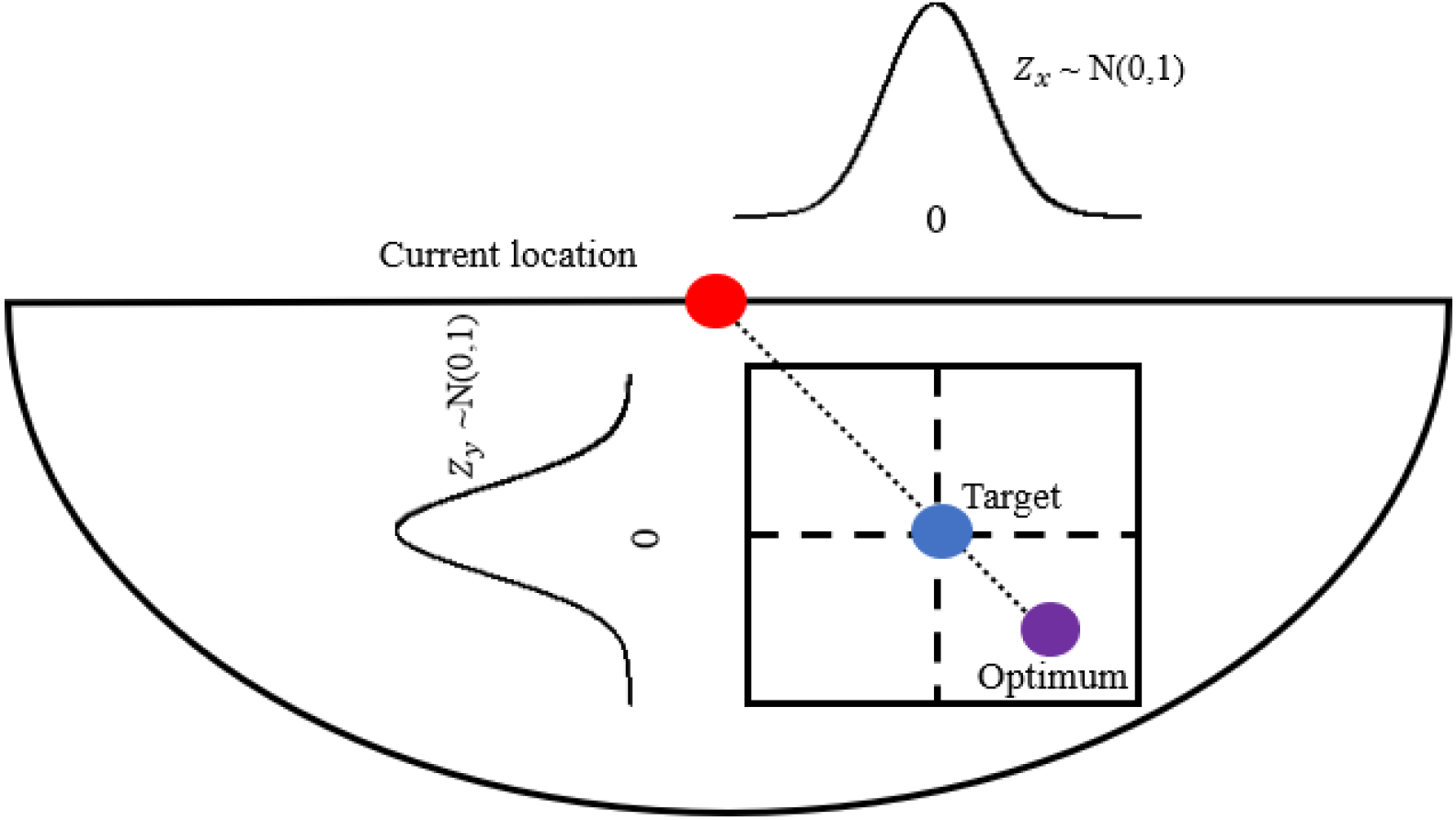
Illustration of large-scale searching specified by the OU model of Eqns. (1) and (2). Agents find an ‘optimum’ location within the semi-circular search region and then a ‘target’ location that lies on the line between the ‘current location’ and the ‘optimum’ location. If there is a tie between multiple potential ‘optimum’ cells, one is randomly selected from the list of tied cells to serve as the optimum. Random variance is added by independently sampling Z_x_ and Z_y_ from a standard normal distribution. Here, *σ* = 1 and ϕ_x_ = ϕ_y_ < 1, where *σ* is the multiplier on the random error and ϕ_x_ and ϕ_y_ are the motivations in the x and y directions, respectively. Bounding box represents the most probable samples from the N(0,1) distribution.

Agents will move toward the selected optimum location. However, if the attraction strength (*ϕ_x_* and *ϕ_y_* in Eqns. (1) and (2)) is less than 1, agents will have a ‘target’ location short of the optimal location. Thus, agents find an ‘optimum’ location within the semi-circular search region and then a ‘target’ location that lies on the line between the ‘current location’ and the ‘optimum’ location. Moreover, agents will move towards this target location with some variance, which is generated by sampling two numbers from the standard normal distribution and multiplying by σ, as specified by the user (In Fig. 3, *σ* is 1). Because the support of the normal distribution consists of all real numbers, large deviations from the ‘target’ point are possible. However, because the normal distribution has low density at the extreme tails, outcomes are most likely to fall within a certain area of the target, as illustrated in Fig. 3. This first step corresponds to the OU model of Eqns. (1) and (2).

The second step is small-scale searching. Here, agents select the ‘best’ of the 8 neighboring cells (queen’s case or Moore neighborhood) after performing step 1, discussed above and in Fig 3. Again, ‘best’ here means the cell with the environmental raster value closest to the agent’s user-defined optimum range. These two steps are then repeated for each timestep until the agent dies or proceeds through all timesteps. Each timestep will use different environmental raster layers. Users may choose to supply a raster stack to simulate changes in environment over time (see for example Section 3.1).

For the `moveSIM` function, agent death occurs when agents fail to achieve suitable environmental raster values for more than a user-specified number of consecutive timesteps. Here, what constitutes a ‘suitable’ cell is determined by the optimum value and an allowable deviation proportion, both also specified by the user. For the `energySIM` function, agent death occurs when energy reaches zero. For both functions, users may choose to disable agent mortality. In the following subsections, we present the differences between `moveSIM` and `energySIM` functions and their underlying algorithms.

### 2.1. Simulation function: moveSIM

The function `moveSIM` runs an OU movement simulation based on environmental conditions provided by the user (e.g., raster), optionally including agent mortality and adjusted motivation according to user-specified parameters. The function operates according to the following algorithm. Here, terms in *italics* are `moveSIM` function arguments (see Table 2).

**Table 2.** List of arguments used in `moveSIM` (M), `energySIM` (E), or `as.species` (S). T: True, F: False, SD: standard deviation. In text, these arguments are presented in *italics.* For a more complete list of argument descriptions, see the abmR documentation.

| Argument | Function | Usage |
| --- | --- | --- |
| replicates | M, E | # of agents to model |
| days | M, E | # of timesteps |
| modeled_species | M, E | Species object from `as.species()` |
| env_rast | M, E | Environmental raster |
| optimum | M | Optimal environmental value |
| optimum_lo | E | Lowest optimum environmental value |
| optimum_hi | E | Highest optimum environmental value |
| dest_x | M,E | Destination Longitude |
| dest_y | M,E | Destination Latitude |
| mot_x | M,E | Motivation (x direction) |
| mot_y | M,E | Motivation (y direction) |
| search_radius | M,E | Radius of semi-circular search region (km) |
| direction | M,E | Movement direction: N, S, E, W, or R (Random) |
| sigma | M,E | Randomness parameter |
| mortality | M,E | Incorporate agent mortality? T or F |
| fail_thresh | M | Deviation from optimum constituting failure |
| n_failures | M | Allowable # of failures before agent death |
| init_energy | E | Initial energy |
| energy_adj | E | Energy gain/penalty vector |
| single_rast | M, E | Using a single-layer raster? T or F |
| write_results | M, E | Save results as a .csv? T or F |
| x | S | Origin longitude |
| y | S | Origin latitude |

The following algorithm applies when the argument *direction* is ‘N’, ‘S’, ‘E’, or ‘W’. For random movement (*direction = ‘R’*) agents simply select a random point from a circle of radius *search_radius* for each timestep (Step 2). Here, let *env_rast* (*x_t+1_, y_t+1_*) be the value of *env_rast* at the point (*x_t+1_, y_t+1_*). The core algorithm shown here assumes that *env_rast* contains no undefined (N/A) grid cells.

1. Specify (x_1_, y_1_) using *x* and *y* contained in *modeled_species*, set failures = 0.
2. For day t in 1:(*days*-1)
  a. Define search area semicircle (radius = *search_radius)* facing *direction* and centered at (*x_t_*, *y_t_*)
  b. Determine (θ_*x*_, θ_*y*_) as location within the search area with *env_rast* value closest to *optimum.*
    i. If (*dest_x, dest_y*) in search area, set (θ_*x*_, θ_*y*_) = (*dest_x, dest_y*)
  c. Large scale searching: find (*x_t+1_, y_t+1_*)_0_ according to (1) and (2).
  d. Small scale searching: set (*x_t+1_, y_t+1_*) as location within eight neighboring cells (queen’s case) of (*x_t+1_, y_t+1_*)_0_ with the value closest to *optimum*. Perform (e)-(f) if *mortality* = True.
  e. If observed *env_rast*(*x_t+1_, y_t+1_*) - *optimum > optimum*fail_thresh*, set failures = failures + 1. If not, set failures = 0.
  f. If failures > *n_failures* agent dies. End loop.
3. Return dataframe with *days* rows and 2 columns movement track data.
4. Repeat (1)-(3) *replicates* times.

### 2.2. Simulation function: energySIM

The function `energySIM` builds on `moveSIM` by allowing for dynamic agent energy levels that are affected by the quality of environmental values achieved. These initial user-defined energy levels then serve as a driver of mortality and movement distance per timestep. The energy thresholds range from 0 to 1 and represent proportion deviations from the optimum. These thresholds are arbitrarily divided into 10 even increments (0.1, 0.2, etc.), but users can change the energy gain or penalty for attaining each of the thresholds. It operates according to the following algorithm. Here, terms in *italics* are `energySIM` function arguments (see Table 2) or calculated variables (e.g., *optimum, energy*).

The following algorithm applies when the argument *direction* is ‘N’, ‘S’, ‘E’, or ‘W’. For random movement (*direction =* ‘ *R*’) agents simply select a random point from a circle of radius *search_radius* for each timestep (Step 3). Here, let *env_rast* (*x_t+1_, y_t+1_*) be the value of *env_rast* at the point(*x_t+1_, y_t+1_*). The core algorithm shown here assumes that *env_rast* contains no undefined (N/A) grid cells.

1. Specify (x_1_, y_1_) using *x* and *y* contained in *modeled_species.*
2. Compute *optimum* as (*optimum_hi - optimum_lo)/2 and* set *energy* = *init_energy*
3. For day t in 1:(*days*-1)
  a. If *mortality* = True, update *search_radius* as *search_radius = search_radius * (energy/init_energy).*
  b. Define search area semicircle(radius = *search_radius*) facing *direction* and centered at (*x_t_*, *y_t_*).
  c. Determine (θ_*x*_, θ_*y*_) as location within the search area with *env_rast* cell value closest to *optimum.*
    i. If (*dest_x, dest_x*) in search area, set (θ_x_, θ_y_) = (*dest_x, dest_y*)
  d. Large scale searching: find(*x_t+1_, y_t+1_*)_0_ according to (1) and (2).
  e. Small scale searching: set (*x_t+1_, y_t+1_*) as location within eight neighboring cells (queen’s case) of (*x_t+1_, y_t+1_*)_0_ with the cell value closest to *optimum*.
  f. Update energy level according to deviation of env_rast (*x_t+1_, y_t+1_*) from optimum.
  g. If *mortality =* True and *energy = 0*, the agent dies. End loop.
4. Return dataframe with *days* rows and 2 columns movement track data.
5. Repeat (1)-(4) *replicates* times.

## 3. EXAMPLE APPLICATIONS

abmR can be used to construct ABM simulations for any desired agent across the globe. In the following examples, we demonstrate the computation capabilities of the `energySIM` function, although a similar workflow also applies for `moveSIM`. In the first example, we show how `energySIM` can be used to compare movements and differential energy allocations of two synthetic populations of 250 agents each (Box 1). In the second example, we replicate the movement pattern of the Painted Bunting (*Passerina ciris*), a well-studied migratory songbird occurring in North America and Mexico.

**Box 1.** R code used for performing the simulations presented in Fig 4. First, `as.species` is called to initialize two populations with different origin locations. Then, `energySIM` is called to perform a movement simulation for each population; parameters that differ between the two simulations are printed in red, functions in blue, and objects in bold. For argument descriptions, see Table 2 and the package manual.

| Population 1 | Population 2 |
| --- | --- |
| `am.pop.1 = as.species(x = -105.7, y = 48.2) sim.move <- energySIM( replicates = 250, days = 14, env_rast = as.raster.stack.ndvi.sep, search_radius = 150, sigma = 0.1, dest_x = 999, dest_y = 999, mot_x = 0.95, mot_y = 0.95, modeled_species = am.pop.1, optimum_lo = 0.2, optimum_hi = 0.5, init_energy = 100, direction = "S", mortality = F, Energy_adj = c(20,10,8,5,2,0,-2,-5,-8,-10,-20), write_results = T, single_rast = F)` | `am.pop.2 = as.species(x = -142.7, y = 63.2) simtwo.move <- energySIM( replicates = 250, days = 14, env_rast = as.raster.stack.ndvi.sep, search_radius = 800, sigma = 0.1, dest_x = 999, dest_y = 999, mot_x = 0.8, mot_y = 0.8, modeled_species = am.pop.2, optimum_lo = 0.6, optimum_hi = 0.8, init_energy = 70, direction = "S", mortality = F, energy_adj = c(20,10,8,5,2,0,-2,-5,-8,-10,-20), write_results = T, single_rast = F)` |

### 3.1. Comparisons between populations

In this example, both synthetic populations are characterized by matching number of replicates and movement timesteps (*days*), equal *σ*, and the same environmental data provided by a Normalized Difference Vegetation Index (NDVI) raster stack containing 14 days of data between September 01-14, 2019 (Vermote, 2019). Both populations had an unspecified destination (indicated with ‘999’ in the arguments *dest_x* and *dest_y*) and were constrained to move on land. However, Population 1 (P1) agents started their movements from a different point (105.7° W; 48.2° N) situated about 2,800 km from the origin of Population 2 (P2) agents (142.7° W; 63.2° N). Additionally, P1 agents had a smaller search radius (150 km) but higher motivation than P2 agents (P1 motivation = 0.95). P1 agents also had different optimum ranges (P1 0.2-0.5; P2 0.6-0.8), and different initial energy units (P1 100; P2 70). These differences in simulation parameterization result in clearly dissimilar movement tracks (Fig. 4).

**Figure 4.**
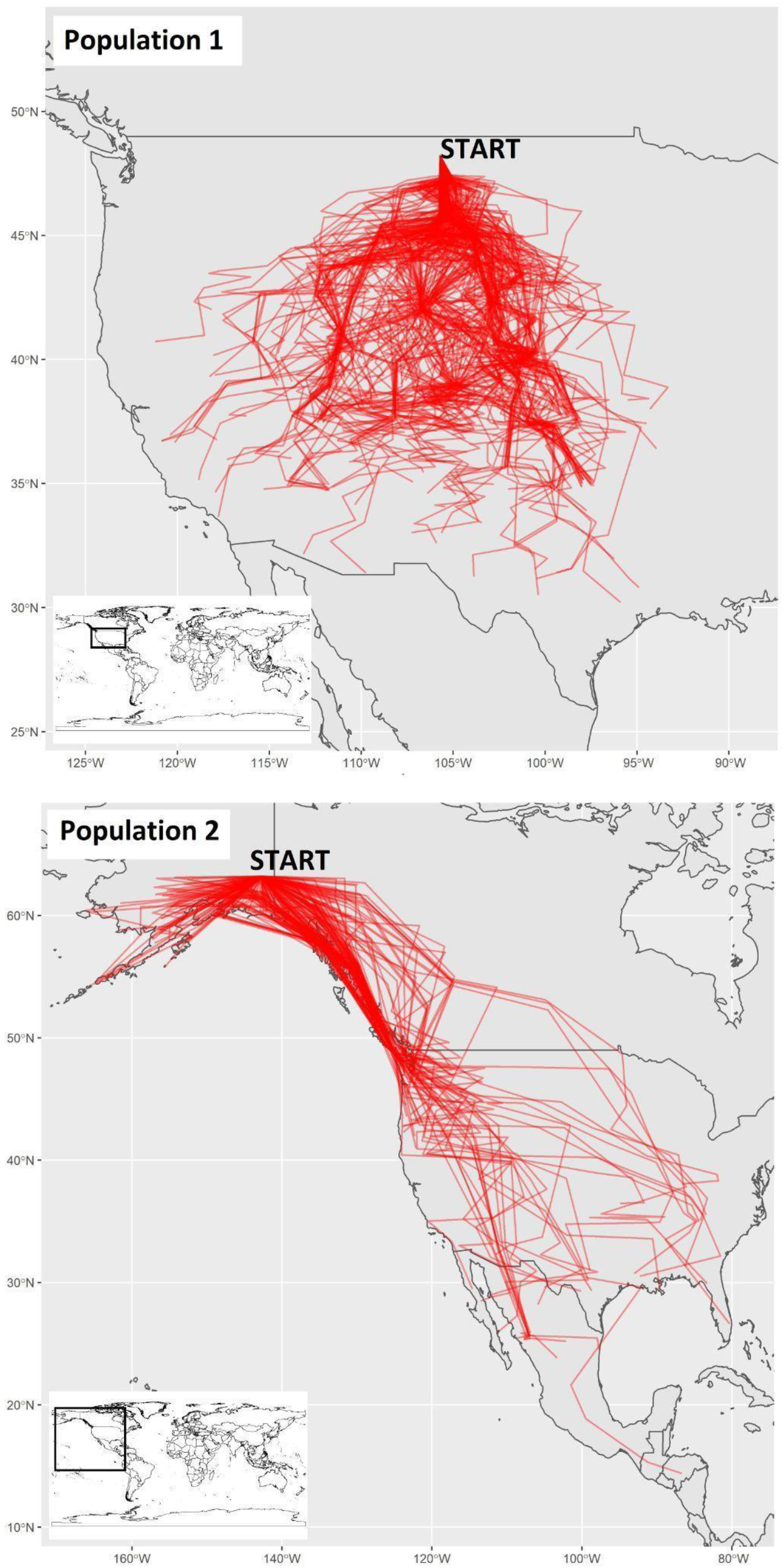
Movement tracks reveal that Population 1 tended to travel through the central United States, while Population 2 traveled mostly throughout western Canada, United States, and Mexico. Overall, Population 1 traveled more distance and exhibited more consistent paths near the origin than did Population 2. The movement tracks are natively produced by abmR. Inset world map provided for geographic reference.

While we can compare the movement tracks visually, Table 3 provides a numerical description of the results. In this simulation, P1 traveled a much smaller average distance (154.6 km) than did P2 (625.5 km). However, P1 traveled more days on average (7.4 days) before stopping than P2 (3.8 days). Additionally, P2 had higher energy consumption than P1; its average remaining energy across all timesteps was 61.2 units compared to 99.7 units for P1. There are several possible reasons for this observed pattern. First, P2 began with a smaller initial energy (70 units) than P1 (100 units). Additionally, P2 had higher optimum NDVI values (0.6 - 0.8), which might have been less abundant and generally more difficult to reach than those of P1 (0.2 - 0.5). Finally, because they began in different places, P1 and P2 agents encountered different raster cells along their journey.

**Table 3.** A numerical comparison of Populations 1 and 2, created by using values across all timesteps for all agents. ‘Day’ summarizes the timestep variable of the movement tracks. ‘Longitude’ and ‘latitude’ summarize the geographical position of agents, while ‘energy’ summarizes agents’ remaining energy. ‘Delta energy’ corresponds to the change (gain or loss) of energy between each timestep, while ‘distance’ refers to the distance traveled between each timestep. This table was produced outside of abmR using raw movement data returned by the package.

|  | variable | mean | sd | median | min | max | range |
| --- | --- | --- | --- | --- | --- | --- | --- |
| Population 1 | day | 7.4 | 4 | 7 | 1 | 14 | 13 |
|  | longitude | -105.5 | 4.1 | -105.2 | -121.1 | -93.3 | 27.8 |
|  | latitude | 41.6 | 3.4 | 41.9 | 30.2 | 47.5 | 17.3 |
|  | energy | 99.7 | 1.3 | 100 | 80 | 100 | 20 |
|  | delta energy | -0.04 | 1.5 | 0 | -20 | 20 | 40 |
|  | distance | 154.6 | 70 | 154.4 | 5.5 | 316.7 | 311.1 |
| Population 2 | day | 3.8 | 3.08 | 3 | 1 | 14 | 13 |
|  | longitude | -126 | 17.3 | -127.1 | -166 | -80.3 | 85.7 |
|  | latitude | 50.2 | 10.5 | 52.2 | 14.4 | 63.1 | 48.7 |
|  | energy | 61.2 | 17.2 | 60 | 0 | 100 | 100 |
|  | delta energy | -2.3 | 9.3 | -5 | -20 | 20 | 40 |
|  | distance | 625.5 | 294 | 649.7 | 5.5 | 1445 | 1439.4 |

Fig. 5 visually compares P1 and P2 movement outputs based on longitude and latitude. This is not a native abmR figure, but rather is produced using the raw data that abmR generates to show the flexible use of the package. In this figure, P1 movements tended to be to the east and south of P2. However, P2 trajectory shows a much wider distribution, with density points extending to the lower values of latitude.

**Figure 5.**
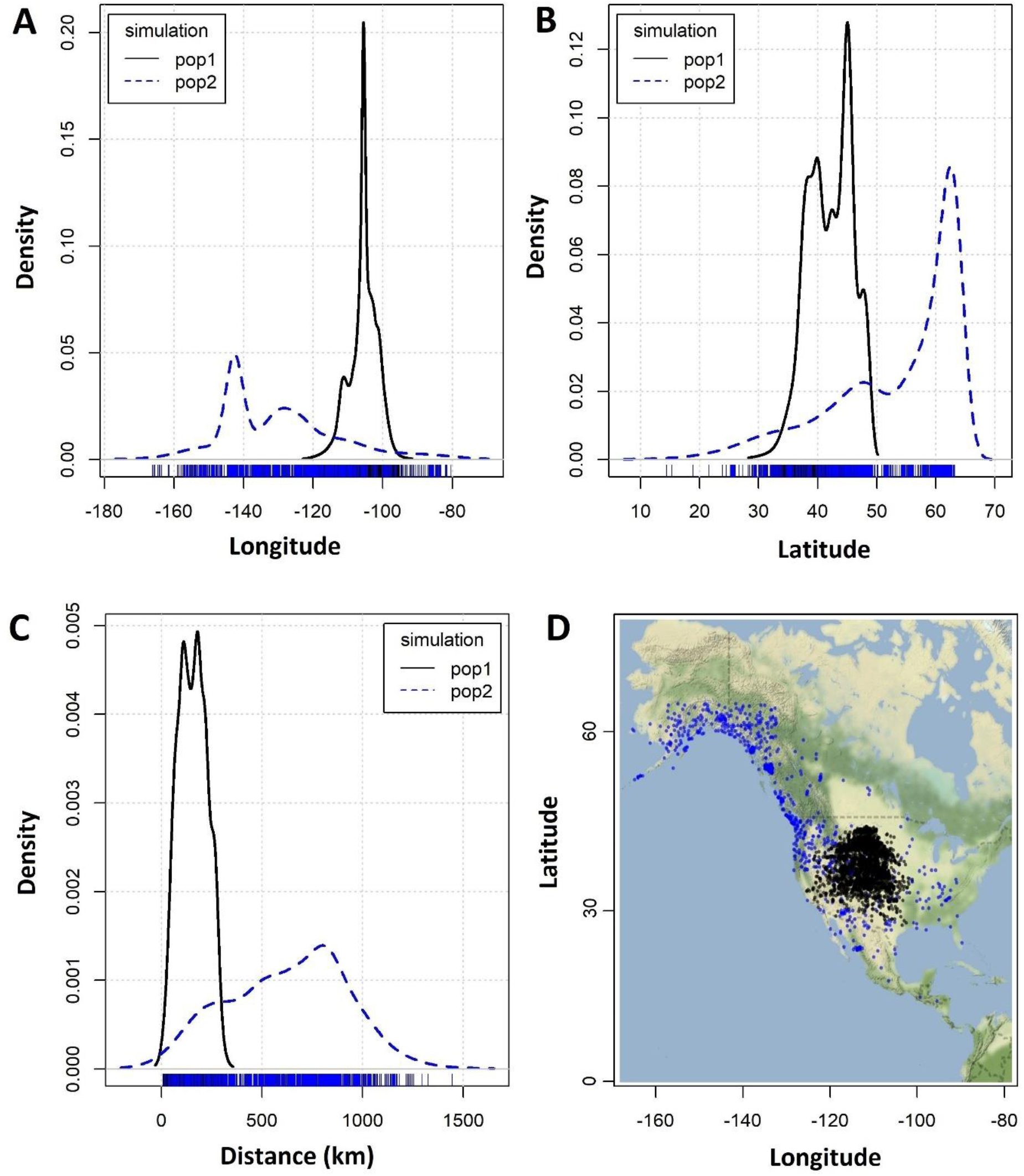
Graphical comparisons of Population 1 and Population 2 movements. Panels A and B show density plots used to individually compare longitude (Panel A) and latitude (Panel B) coordinates attained by agents from each population. Panel C compares the distance traveled between each timestep, while Panel D shows geographical position for all agents in each population across all timesteps.

Finally, Fig. 6 provides a density surface plot for P2 describing agent energy gains (blue) and losses (red) across the landscape. This surface was created using the inverse distance weighted interpolation (IDW) function from the R package ‘gstat’ (Pebesma, 2004). IDW interpolates grid cell values across a surface using a linear combination of observed (sample) points. When interpolating a cell value, the value of the sample points closer to that cell carry a higher weight, while sample points further from that cell carry smaller weight. IDW is discussed in more detail in Wong (2017). The results from Fig. 6 match well with what we observe in Fig. 4. Movement tracks for P2 tend to follow the blue (energy gain) regions.

**Figure 6.**
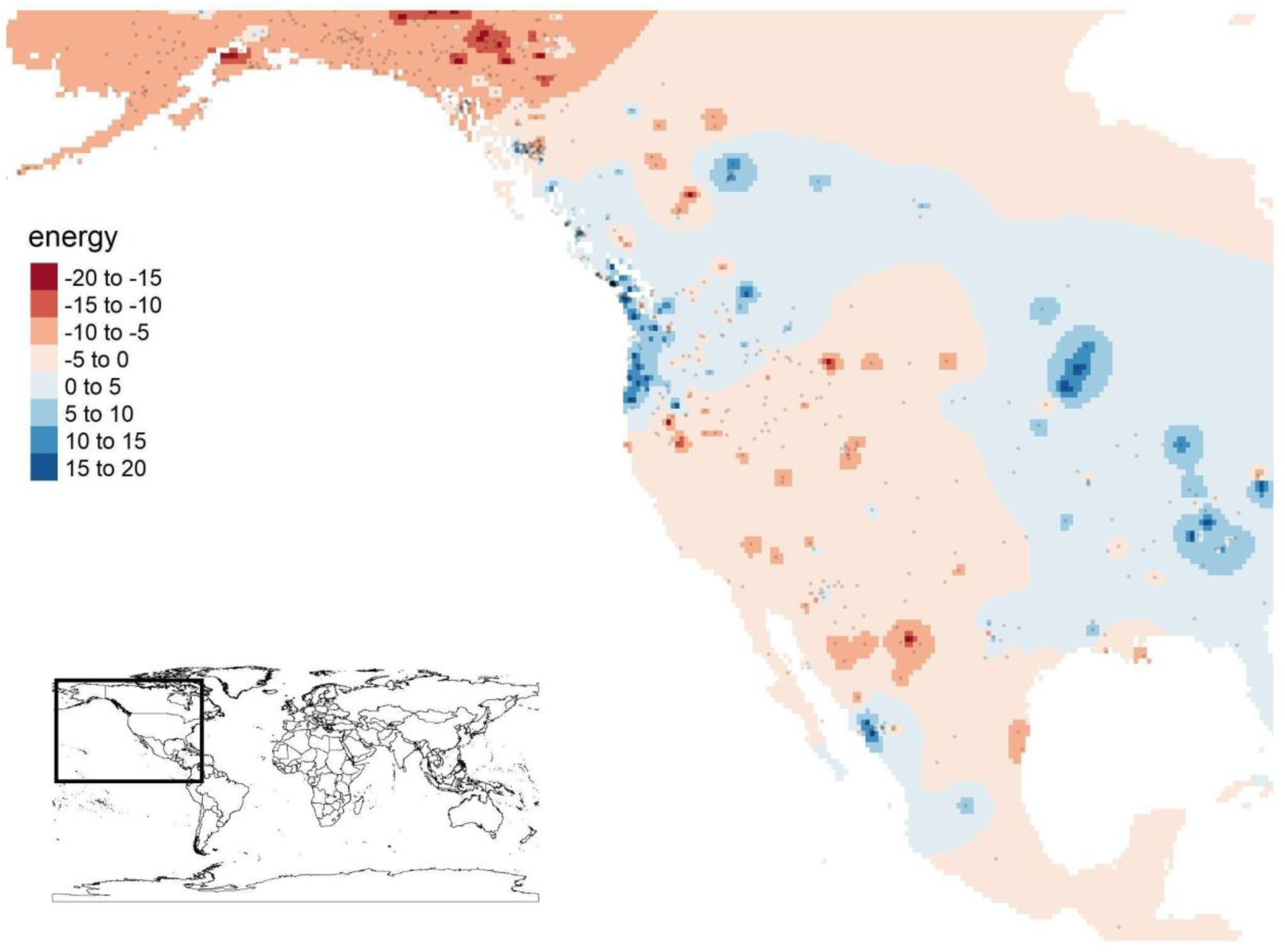
Energy gradient plot of Population 2 by timestep. Areas in red reflect energy loss (less suitable environmental values) while areas in blue reflect energy gain (better environmental values). This plot is produced directly by energyVIZ. Inset world map added for geographic reference.

### 3.2. Migration of the Painted Bunting

The Painted Bunting is a migratory songbird that has been intensively studied over the past two decades due to its steeply declining population trend in the United States (U.S.) and its complex molt-migratory behavior (Thompson 1991). This species has been the focus of pioneering light-level geolocator tags that revealed westward post-breeding movements from the southern U.S. throughout Mexico (Contina et al. 2013), a system for genetic analysis of migratory population connectivity (Battey et al. 2018; Contina et al. 2019) and for candidate genes studies related to the migratory behavior (Contina et al. 2018). Moreover, Bridge et al. (2016) modeled the post-breeding movements of a Painted Bunting population departing Oklahoma (U.S.) through an ABM approach and investigated the association between large-scale vegetation productivity changes and the molt-migratory behavior. Therefore, this species provides a well-studied migratory system with ample behavioral and population ecology knowledge against which abmR movement predictions can be tested and compared.

We built a basic ABM movement simulation for the Painted Bunting in abmR by using the same key parameters adopted by Bridge et al. (2016) and predicted that a westward migratory movement would emerge throughout the southern U.S. and the coastal regions of Mexico in late summer. Bridge et al. (2016) found that Painted Bunting agents beginning their migratory journey in Oklahoma in late August and September tend to avoid a direct southern migration and show a clear pattern towards southwestern movements targeting high primary productivity areas in northern Mexico and Sinaloa (northwestern Mexico). In our simulation, we calibrated our model by following the movement parameters adopted by Bridge et al. (2016). We used the identical model start location (−98.8° W; 34.8° N), a similar subset of 14 vegetation index raster files (NDVI; Vermote, 2019) representing primary productivity condition during the first two weeks of September 2011, and no predefined destination coordinates (indicated with ‘999’ in the arguments *dest_x* and *dest_y*). For the full set of model parameters, see suppl. material S2. While we note that Bridge et al. (2016) used enhanced vegetation index (EVI) as opposed to NDVI, a multiyear timeframe (2010-2013), and a complex series of model parametrization implemented in ArcGIS, our simulation offers a basic but useful illustration of the predictive capabilities of abmR.

The outcome of 12 abmR Painted Bunting simulations revealed a movement pattern consistent with our predictions based on the results presented by Bridge et al. (2016). Most agents in our simulation (9 out of 12) showed a southwestern movement towards Sinaloa (Mexico) in mid-September (Fig. 7), where migratory agents utilize a bloom in primary productivity due to the Monsoonal precipitations (Rohwer et al. 2005). Three agents showed a southeastern movement, following vegetation changes along southeastern U.S. and the Gulf of Mexico. This result is also in line with sporadic but nonetheless documented variations in migratory strategies in the Painted Bunting where some individuals that breed in south central U.S. (e.g., western Oklahoma and Arkansas) show southeastern movements in late summer (Contina et al. 2013).

**Figure 7.**
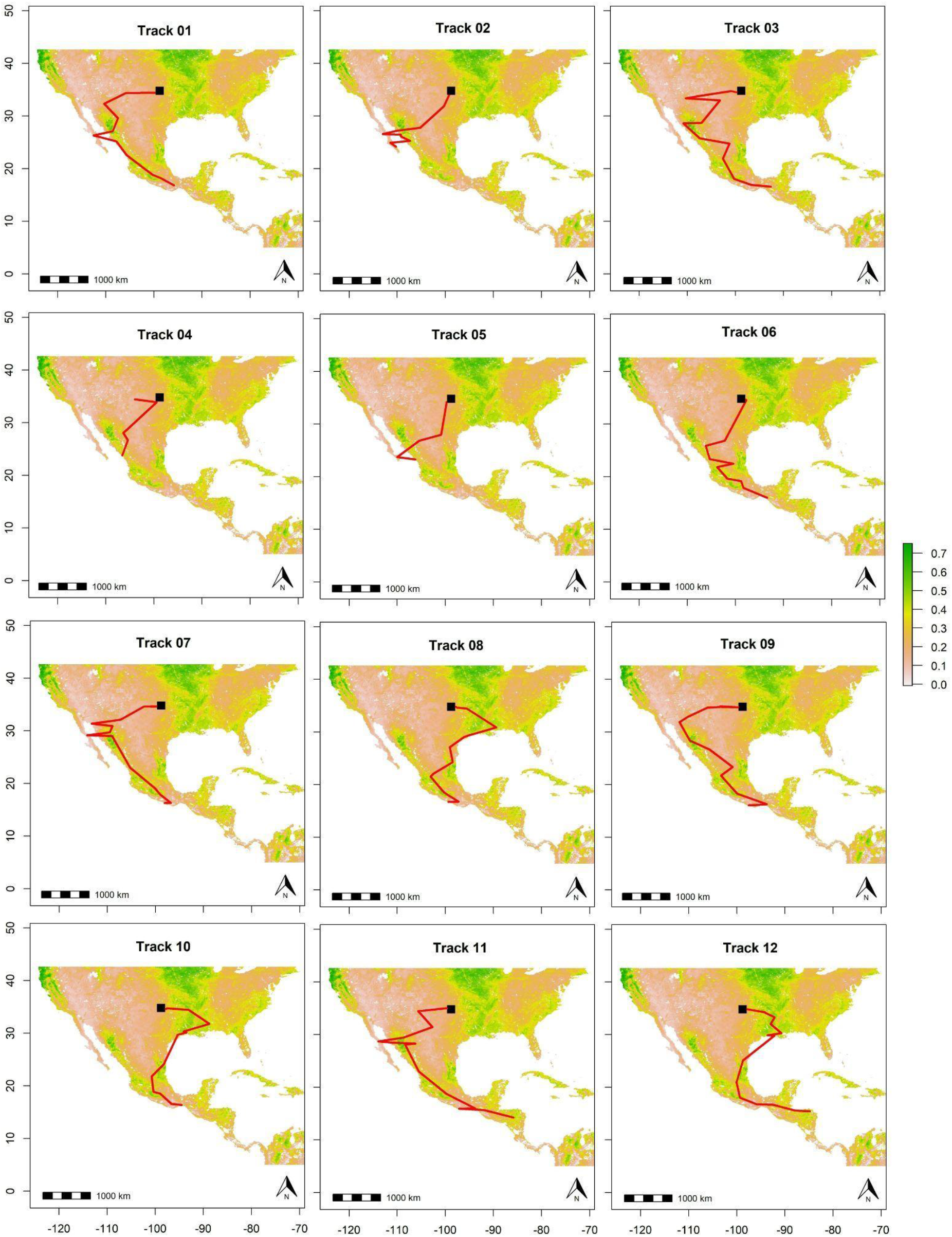
Outcome of 12 abmR simulations showing a frequent southwestern migration from the breeding ground in Oklahoma (U.S.) towards wintering grounds in Sinaloa (Mexico) and southern Mexico. The NDVI map in the background is an average of the raster stack object used in abmR which contained 14 raster layers ranging from September 1 to September 14, 2011.

## 4. CONCLUSIONS AND FUTURE WORK

abmR provides a novel and efficient programming platform for simulating large-scale movements of species across taxa. We ran most of the initial test simulations on a local machine equipped with an Intel® Core™ i7-5500U CPU – 2.40GHz and 8 GB of RAM and obtained results for 100-1000 agents within minutes. The novelty of the software includes the capability of concurrently modeling agent movement trajectories and energy budget. This feature enables a broader exploration of the ecological constraints that shape animal dispersal and/or migration. Moreover, abmR built-in arguments, such as *fail_thresh, n_failures*, and *energy_adj*, provide additional flexibility when evaluating mortality scenarios that depend on baseline environmental conditions and energy requirement during prolonged movement bouts (see Table 2 for a full list of arguments affecting mortality).

Over the last decades, spatially explicit simulations, pattern-oriented modelling, approximate Bayesian computing, and agent-based models have become more popular in ecological and evolutionary studies (Railsback et al., 2006; DeAngelis & Grimm, 2014; van der Vaart et al. 2015; Gallagher et al. 2021). Analytical platforms, such as InSTREAM, a simulation model approach designed to understand how stream and river salmonid populations respond to habitat alteration (Railsback et al., 2009), or ALMaSS, a predictive modeling tool for answering environmental policy questions regarding the effect of changing landscape structure on threatened animal species (Topping et al., 2003), allow investigation of specific ecological systems using ABM. On the other hand, many programming languages such as Netlogo, R, or Python are widely used to develop custom and more flexible models that can be adapted to address complex ecological or evolutionary research scenarios (Lustig et al., 2019; Chubaty & McIntire, 2021). However, the use of a programming language to develop a flexible ABM from scratch has two important drawbacks. First, it requires advanced programming skills. Second, its reproducibility can be compromised by the idiosyncrasies of the simulation algorithm written by the user. These idiosyncrasies, especially if not well documented, can make it difficult or even impossible for other researchers to replicate findings or adapt code to suit their modeling scenarios. abmR provides a novel framework to perform complex movement simulations through standardized functions and arguments that facilitate model annotation and reproducibility while providing publication-ready visualizations at the end of each run.

While we developed and tested abmR as a movement and energy budget simulation tool, its core software functionalities can be adapted to explore other processes such as disease outbreak scenarios (Dougherty et al., 2018). As an example, pathogen vector movement can be easily simulated within abmR, allowing the study of areas of confluence where disease transmission is more probable (Manore et al., 2015). Moreover, potential future updates will include the ability to specify multiple raster stacks of different movement predictors and interactions between agents. In abmR, each simulation output can be used as the input for the next movement model. However, the option of computing agent interactions affecting movement patterns within the same simulation run is currently missing. This is a clear area of further package development. Additionally, other code expansions might be useful to study plant seed dispersal, density-dependent scenarios, and altitudinal movements.

## Acknowledgements

We thank three anonymous reviewers who provided invaluable feedback on an earlier draft. abmR relies heavily on a large set of R packages and we thank the R community for providing support and open-source code. We thank Jeff Kelly, Xiangming Xiao, and the Center for Earth Observation and Modeling at the University of Oklahoma for supporting the research effort.

## Author Contributions

B.G. led the development of the package and wrote the manuscript; J.F.L. contributed to the package, writing, and critical review of the manuscript; A.C. conceived the manuscript, led its writing, contributed to the package, and performed package testing.

## Data Availability

abmR is available on CRAN and the simulation results for Populations 1 and 2 are available for download from Zenodo. The environmental raster data set used in the examples is available at https://www.ncei.noaa.gov.

