## Supplementary Material S1 for "abmR: an R package for agent-based model analysis of large-scale movements across taxa"

Gochanour et al.

moveSIM algorithm:

1. Specify (x_1_, y_1_) using *x* and *y* contained in *modeled_species*.
2. Set failures = 0
3. For day t in 1:(*days*-1)
   a. Create a search area defined as a semicircle of radius *search_radius* facing *direction* and centered at ($x_{t}$, $y_{t}$)
   b. Determine ($\theta_{x}$, $\theta_{y})$ as location within the search area with *env_rast* cell value closest to *optimum.* Draw a random sample (n = 1) in case of ties.
      i. If (*dest_x*, *dest_y*) in search area, set ($\theta_{x}$, $\theta_{y})$ = (*dest_x*, *dest_y*)
   c. Large scale searching: determine $(x_{t+1}$, $y_{t+1}$)_0_ according to (1) and (2).
   d. Small scale searching: determine $(x_{t+1}$, $y_{t+1}$) by selecting location within eight neighboring cells (queen’s case) of $(x_{t+1}$, $y_{t+1}$)_0_ with the cell value closest to *optimum*. Draw a random sample (n = 1) in case of ties.

Perform (e)-(f) if *mortality* = True.

e. If observed *env_rast*$(x_{t+1}$,$y_{t+1}$) - *optimum > optimum*fail_thresh,* set failures = failures + 1. If not, set failures = 0.
f. If failures > *n_failures*, the agent dies. Set $(x_{t+1}$,$y_{t+1}$) through $(x_{days}$,$y_{days}$) as N/A and end loop.

4. Return dataframe with *days* rows and 2 columns movement track data.
5. Repeat (3)-(5) *replicates* times.

energySIM algorithm:

1. Specify (x_1_,y_1_) using *x* and *y* contained in *modeled_species*.
2. Compute *optimum* as (*optimum_hi - optimum_lo)/2.*
3. Set *energy* = *init_energy*
4. For day t in 1:(*days*-1)
   a. If *mortality* = True, update *search_radius* as *search_radius = search_radius * (energy/init_energy).*
   b. Create a search area defined as a semicircle of radius *search_radius* facing *direction* and centered at ($x_{t}$, $y_{t}$).
   c. Determine ($\theta_{x}$, $\theta_{y})$ as location within the search area with *env_rast* cell value closest to *optimum.* Draw a random sample (n = 1) in case of ties.
      i. If (*dest_x*, *dest_y*) in search area, set ($\theta_{x}$, $\theta_{y})$ = (*dest_x*, *dest_y*)
   d. Large scale searching: determine $(x_{t+1}$,$y_{t+1}$)_0_ according to (1) and (2).
   e. Small scale searching: determine $(x_{t+1}$,$y_{t+1}$) by selecting location within eight neighboring cells (queen’s case) of $(x_{t+1}$,$y_{t+1}$)_0_ with the cell value closest to *optimum*. Draw a random sample (n = 1) in case of ties.
   f. If *optimum_lo* < *env_rast*$(x_{t+1}$,$y_{t+1}$) < *optimum_hi* update *energy = energy+energy_adj*[1].
   g. Else, compute $\Delta=$*env_rast*$(x_{t+1}$,$y_{t+1}$) - *optimum.* And update energy in the following way:

If $\Delta$< 0.1 * *optimum*, then *energy = energy+energy_adj*[2]

If 0.1 * *optimum<* $\Delta$< 0.2 * *optimum*, then *energy = energy+energy_adj*[3]

If 0.2 * *optimum<* $\Delta$< 0.3 * *optimum*, then *energy = energy+energy_adj*[4]

If 0.3 * *optimum<* $\Delta$< 0.4 * *optimum*, then *energy = energy+energy_adj*[5]

If 0.4 * *optimum<* $\Delta$< 0.5 * *optimum*, then *energy = energy+energy_adj*[6]

If 0.5 * *optimum<* $\Delta$< 0.6 * *optimum*, then *energy = energy+energy_adj*[7]

If 0.6 * *optimum<* $\Delta$< 0.7 * *optimum*, then *energy = energy+energy_adj*[8]

If 0.7 * *optimum<* $\Delta$< 0.8 * *optimum*, then *energy = energy+energy_adj*[9]

If 0.8 * *optimum<* $\Delta$< 0.9 * *optimum*, then *energy = energy+energy_adj*[10]

If $\Delta$> 0.9 * *optimum*, then *energy = energy+energy_adj*[11]

h. If *mortality =* True and *energy = 0*, the agent dies. Set $(x_{t+1}$,$y_{t+1}$) through $(x_{days}$,$y_{days}$) as N/A and end loop.

5. Return dataframe with *days* rows and 2 columns movement track data.
6. Repeat (3)-(6) *replicates* times.
