## Supplementary Material S2 for "abmR: an R package for agent-based model analysis of large-scale movements across taxa"

Gochanour et al.

ABM movement simulation for the Painted Bunting in abmR

#----------------------------------------------------------------

setwd("E:/abmR_Contina/PABU_example")

am.pop.1 = as.species(x=-98.8, y=34.8)

sim.move <- energySIM(

replicates = 1,

days = 14,

env_rast = as.raster.stack.ndvi,

search_radius = 600,

sigma = 0.3,

dest_x = 999,

dest_y = 999,

mot_x = 0.7,

mot_y = 0.7,

modeled_species = am.pop.1,

optimum_lo = 0.6,

optimum_hi = 0.8,

init_energy = 100,

direction = "S",

mortality = T,

energy_adj=c(20,10,8,5,2,0,-2,-5,-8,-10,-20),

write_results = T,

single_rast = F)

energyVIZ(sim.move)

#=================================================================

jpeg(file = "./sim.move.12.jpeg", width = 8, height = 8, units = 'in', res =400)

energyVIZ(sim.move)

dev.off()
